# Concomitant Vascular Calsequestrin 2 overexpression and leukocytes transmigration in a rat model of skeletal muscle traumatic inflammatory injury

**DOI:** 10.1101/2022.09.29.510069

**Authors:** Noureddine Ben Khalaf, Dalal Al-Mehatab, Dahmani M. Fathallah

**Author notes:** **Corresponding author:** Prof. Dahmani M. Fathallah, Department of Life Sciences, Health Biotechnology Program, College of Graduate Studies, Arabian Gulf University, Road 2904 Building 293 Manama, 329, Bahrain.

## Abstract

The network of molecular mediators involved in the transmigration of leukocytes to inflamed tissues has been expanding with the identification of new molecules involved in the inflammatory response. We have previously shown using a rat model that Protein Disulfide Isomerase PDIA4 (ERP72) is involved in the inflammatory response to skeletal muscle traumatic injury. In this paper, we report observations suggesting that calsequestrin 2 (CASQ2), another member of the thioredoxin/PDI family, might contribute to the inflammatory response that leads to adhesion and transmigration of leukocytes into injured skeletal muscle. Indeed, real time PCR assay showed that the expression level CASQ2 is significantly enhanced [p<0.01] in the mechanically injured muscle. Immunohistological and immunofluorescence analysis showed that CASQ2 but not CASQ1 is expressed in the vessels present in the muscle-injured area. In addition, CASQ2 overexpression and PMN transmigration to the inflamed muscle injury are concomitant. This overexpression occurs most likely in the smooth muscle of the inflamed vessel. These observations together with the known Ca^2+^ buffering role of CASQ2 suggest that the protein contributes to the inflammatory process by providing the Ca^2+^ needed to activate the inflammasome and the leukocytes adhesion molecules which enhances transmigration of inflammatory cells into the injured muscle.

## Introduction

The inflammatory response that occurs at the site of a tissue injury involves a variety of cell types and the activation of a large network of mediators acting in a coordinated cascade resulting in the recruitment of leukocytes into the inflamed tissue [1]. In addition to the soluble mediators, this network includes the inflammasome and a large number of cell adhesion molecules (CAMs) expressed on the surfaces of leukocytes and vascular endothelial cells [2]. The inflammasome or NLRP3 (NOD-, LRR- and pyrin domain-containing protein 3) complex, is a molecular complex that senses cellular stress in injured tissues and triggers the inflammatory process [3-6]. Ca^2+^ signaling has been shown to play a critical role in the activation of the NLRP3 inflammasome [7]. Indeed, Endoplasmic Reticulum (ER) release and influx of extracellular Ca^2+^ are both required to activate the NLRP3 inflammasome [8]. On another hand, the activation of inflammatory cells such as monocytes and neutrophils and the regulation of their adhesion are also dependent on calcium mobilization [9-11]. Indeed, the activation of the inflammatory molecular mediators such as the leukocytes adhesion molecules integrins, ICAMs (intercellular adhesion molecules) [12, 13] and L-selectin (CD62L) [14] in physiological as well as inflammatory pathological states have an absolute requirement for calcium ions. Calcium signaling is also involved in the endothelial selectin, E-selectin and the vascular cell adhesion molecule 1 (VCAM1) adhesive functions [14, 15].

In addition, a role for thioredoxin family members e.g. Protein Disulfide Isomerase A1 (PDIA1) and Protein Disulfide Isomerase A4 (PDIA4) has been demonstrated in previous studies, notably in vascular endothelial cells where leukocytes undergo the transmigration process initiated by their adhesion to activated endothelial cells. Indeed, PDIA1was shown to be involved in the refolding-dependent activation of key adhesion molecules such integrins [16]. Furthermore, we previously demonstrated, using a rat model of inflammatory skeletal muscle traumatic injury, that PDIA4 (Erp72) is upregulated in the endothelium of blood vessels post-injury, and that anti-Erp72 antibodies inhibit adhesion of neutrophils to TNFα-activated Human umbilical vein endothelial cells (HUVECs) [17]. Interestingly, the thioredoxin family includes Calsequestrin (CASQ) a protein of the sarcoplasmic reticulum (SR), first discovered in rabbit skeletal muscle tissue in 1971 [18]. Two isoforms of the protein are currently known, CASQ1 and CASQ2, the first was well characterized as a major Ca^2+^-buffering protein in the sarcoplasmic reticulum [19, 20]. CASQ1 helps maintaining the amount of Ca^2+^ and regulates its release from the SR to the cytosol, as well as its entry from extracellular environment. Resolution of CASQ1 structure at a 2.02 Å revealed the presence of three similar domains (I, II and III) with a thioredoxin-like topology and a high capacity of calcium binding [21]. Each domain forms a disk-like shape with a fold of a five-stranded β-sheet in the core flanked by four α-helices [18]. Meanwhile, the CASQ2 solved structure showed that the two isomers, share a common fold pattern. The difference is located at the C-terminus region, where CASQ2 presents a highly extended acidic C-terminus region [18]. To our best knowledge, the function of thioredoxin domains in CASQs is poorly understood, and no link to metabolic, oxidative, signaling or protein folding pathway was established for these proteins, despite the presence of these functional domains. Interestingly, CASQ1 was reported to be involved in several human skeletal muscle diseases, although knock out mice are viable, fertile, and present normal muscular performance, with only a mild atrophy observed. Mutations in the CASQ1 gene have been associated with a modulation of Ca^2+^-dependent tubular aggregation and SOCE-mediated Ca^2+^ homeostasis, inducing symptoms similar of those of malignant hyperthermia [22-28].

In the current study, considering the important role of Ca2+ in the inflammatory process and that CASQs do not display all the structural features of the thioredoxin family previously shown to be involved in this process, we used the rat model of skeletal muscle inflammatory injury [29] to investigate the expression profile of CASQ1 and CASQ2 proteins during the inflammatory process which might indicate a potential link with Ca^2+^ buffering during this process.

## Material and Methods

### Bioinformatic annotations

GenBank database (NCBI) was used to retrieve coding sequences for CASQ1 and CASQ2 proteins. For protein annotation, conserved domains were predicted using Pfam (https://pfam.xfam.org/) and SMART (http://smart.embl-heidelberg.de/) servers. Transmembrane domain was predicted using TMHMM server (http://www.cbs.dtu.dk/services/TMHMM/). Signal Peptide was predicted using SignalP server (http://www.cbs.dtu.dk/services/SignalP-3.0/). Sequence alignment was performed using Clustal Omega (https://www.ebi.ac.uk/Tools/msa/clustalo/). Crsytal structure of CASQ1 (PDB id: 3UOM) and CASQ2 (PDB id: 2VAF) were downloaded from RCSB database (https://www.rcsb.org/), and Pymol (Schrodinger Inc., CA, USA) was used for structure visualization and alignment.

### Primer Design

*Rattus norvegicus* genes: *CASQ1* (Calsequestrin 1*), CASQ2* (Calsequestrin 2), *ICAM1* (intercellular adhesion molecule 1), *VCAM1*(vascular cell adhesion molecule 1), and *SELE* (Selectin E) mRNA sequences were downloaded from Uniprot server (https://www.uniprot.org) and specific primers were designed using Primer3 web server (http://bioinfo.ut.ee/primer3-0.4.0/) default parameters and their specificity assessed using Blast (https://blast.ncbi.nlm.nih.gov) for cross-priming. Primers used for real-time PCR (Table 1) were purchased from Biolegio (Biolegio, Nijmegen, The Netherlands) in lyophilized form. All primers were synthesized at a 40μM scale and purified by HPLC. The primers were re-suspended in Nuclease free water at a concentration of 100μM.

**Table 1.**
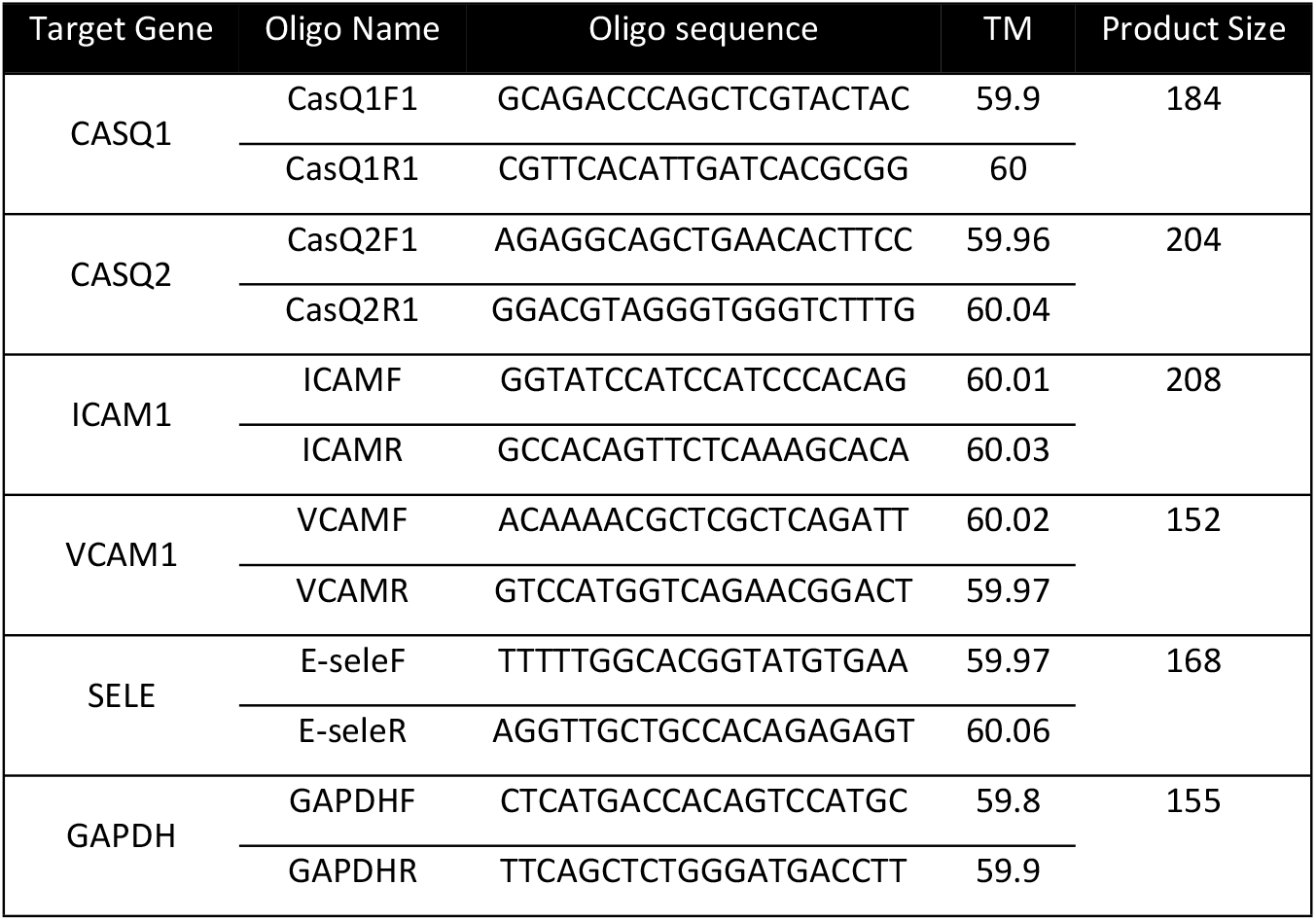
Primer List.

### Animals

Thirty Inbred Wistar female rats weighing 200–220 g were used to develop the muscle injury model, following institutional guidelines and in conformity with the international standards recommended for animal experimentation. Five groups of five rats were used for the monitoring of gene expression and Immunohistochemistry experiments post-injury. One animal group did not undergo muscle injury and was used as an untreated control group.

### Rat skeletal muscle injury model

All animal experimental protocols were approved by the Research and Ethics committee at the Arabian Gulf University under the Research Grant LS_NB_18. All methods were carried out in accordance with the committee’s relevant guidelines and regulation. Rat skeletal muscle injury was performed as described previously [29]. Animals were anesthetized intraperitoneally using a mixture of Ketamine at 90 mg/kg and Xylazine at 10 mg/kg (Unimedical LTD, GB). Anesthesia induction was confirmed by pedal reflex. The muscles in both limbs were punctured using a 20-gauge needle mounted on a manual leather-puncturing device to create a hematoma as previously described [29]. The rats were euthanized using CO2 at a displacement at a flow rate of 50% at different time-points varying from 30 min to 4 hours post-injury. A surgical procedure was then used to extract vessels by dissection of the rats’ posterior legs in the injured and non-injured area, vessels were further dissected under microscope to remove all irrelevant tissues. Vessels were then stored in Trizol (Invitrogen, Carlsbad, CA, USA) reagent for RNA extraction. For the Immunohistochemistry study, wounded muscles were resected, formalin-fixed and paraffin-embedded; 4–5 μm thick sections were then cut and stained with hematoxylin and eosin for light microscopy examination. Histological observations were performed on a Zeiss axioskop light microscope (Zeiss, Oberkochen, Germany) using an eyepiece graticule grid. All rats’ samples were considered for the study.

### Gene expression analysis

Total mRNA was extracted from homogenized tissues using the Rneasy RNA extraction kit (QIAGEN) and reverse transcribed using the ProtoScript® First Strand cDNA Synthesis Kit (NEB, UK) according to the manufacturer’s instructions. Primer sets for real-time PCR (Table 1) were used to amplify target regions from genomic DNA and cDNA as templates by the GoTaq DNA polymerase (Promega, WI, USA). For real-time PCR experiments, PowerUp Sybr Green Master Mix (Thermo Fisher Scientific, MA, USA) was used to measure *CASQ1, CASQ2, ICAM1, VCAM1*, and *SELE* gene expression level at 30-, 90- and 120-min post-injury and fluorescence was monitored for 40 cycles on a 7500 fast real-time PCR system (Thermo Fisher Scientific, MA, USA). Experiments were run in triplicates on all groups and results were expressed as the fold change (2-^ΔΔct^ method) using GAPDH (Glyceraldehyde 3-phosphate dehydrogenase) as a housekeeping gene and untreated control group as reference.

### Multiplex Immunohistochemistry

Immunohistochemistry staining was performed on the Ventana Discovery Ultra Chromogenic AmpHQ automated immunostainer (Ventana Medical Systems, AZ, USA) to investigate CASQ1 and CASQ2 protein expression at 4 hours post-injury. The multiplex technology uses the sequential application of unmodified primary antibodies with specific heat deactivation steps in between that does not affect epitope in the tissue [30]. In a sequential staining procedure, deactivation of the primary antibody and secondary antibody-HRP/AP bound to the first biomarker, before the application of subsequent biomarker(s), is critical to reducing cross-reactivity and facilitating downstream image analysis [31]. The Cell Conditioning2 buffer (CC2, #950-123, Ventana, AZ, USA) was used for deactivation of the bound primary antibody and secondary antibody-HRP while maintaining the integrity of the tissue morphology and the subsequent epitopes [30]. Deparaffinization and on-board antigen retrieval were performed for 64 min at 95°C with the CC1 reagent (#950-500, Ventana AZ, USA). Blocking Buffer (Invitrogen Cat# 00-4952-54, Thermo Fisher Scientific, MA, USA) was used for section blocking. Slides were colored using Ventana reagents (Ventana, AZ, USA) except as noted, according to the manufacturer’s instructions. Pre-diluted primary antibody, anti-CASQ1 (1:10000, Abcam # ab185220, Abcam, Cambridge, UK) or anti-CASQ2 (1:10000, Abcam #ab96387, Abcam, Cambridge, UK), was applied first, followed by Omap anti-Ms HRP (Ventana Discovery #760-4310, Ventana, AZ, USA), followed by Teal-HRP kit (Ventana Discovery #760-247, Ventana, AZ, USA) or Purple-HRP kit (Ventana Discovery Purple Kit #760-229, Ventana, AZ, USA) respectively [30]. Finally, the slides were counterstained with hematoxylin and bluing reagent according to the manufacturer’s recommendations. Images acquisition and quantitative analysis were performed using HALO™ Image Analysis Software (PerkinElmer, Inc., MA, USA).

### Triplex Immunofluorescence

For Triplex immunofluorescence, we performed deparaffinization and on-board antigen retrieval for 64 min at 95°C with the CC1 reagent (#950-500, Ventana, AZ, USA). Blocking Buffer (Invitrogen Cat# 00-4952-54) was used for section blocking. Slides were colored using Ventana reagents (Ventana, AZ, USA) according to the manufacturer’s instructions. A Triplex protocol was used, three pre-diluted primary antibodies were sequentially applied in the following order using the indicated chromogenic detection: anti-CASQ1 (1:10000, Abcam # ab185220, Abcam, Cambridge, UK) with Omap anti-Ms HRP (Ventana Discovery #760-4310, Ventana, AZ, USA) and Discovery Cy5 Kit (Ventana Discovery #60-238, Ventana, AZ, USA), followed by anti-CASQ2 (1:10000, Abcam #ab96387, Abcam, Cambridge, UK) with Omap anti-Ms HRP (Ventana Discovery #760-4310, Ventana, AZ, USA) and Ventana Discovery DCC kit #760-240; rabbit anti-CD34 (1:400, Abcam #ab81289) with Omap anti-Rb HRP (Ventana Discovery #760-4310, Ventana, AZ, USA)) and Discovery Rhodamine kit (Ventana Discovery # 760-233, Ventana, AZ, USA). Sections were then stained with DAPI and mounted using Vectashield mounting medium (Vector Laboratories, CA, USA). Images were acquired on Zeiss Axio Observer Z1 Inverted Fluorescence Microscope (Zeiss, Oberkochen, Germany). Images acquisitions were performed using HALO™ Image Analysis Software (PerkinElmer, Inc., MA, USA).

### Statistical Analysis

Experiments of Gene expression analysis were run in triplicates. The two-tailed t-test for independent samples was used to calculate the level of significance (*p*<0.05, *p*<0.01, and *p*<0.001) between any condition compared to the control for all experiments. All calculations were performed using SPSS statistics software version 19 (www.ibm.com).

## Results

### Bioinformatic analysis

CASQ1 and 2 protein sequences were retrieved from the Uniprot database. SMART protein analysis showed the presence of Thioredoxin6 PFAM domain on both proteins, with amino acid position ranging from 186 to 379 for CASQ1, and 171 to 364 for CASQ2, at a significant e-value (2.8e-21and 1.3e-2), respectively. TMHHM analysis showed no transmembrane domain in both proteins. However, Signal P Hidden Markov Models (HMM) analysis, predicted a signal peptide for both CASQ1 And CASQ2, at a high probability (0.999 and 0.994 respectively), with amino acid position ranging from 1 to 34 for CASQ1 and 1 to 19 for CASQ2. Clustal Omega alignment showed 66.4% of identity between CASQ1 and 2 proteins’ sequences as shown by Figure 1. CASQ1 and CASQ2 proteins 3D models’ superimposition shows a conserved structure between the two proteins, except for the extended C-terminal region in CASQ2 (Figure 2) which likely has no major impact on the proteins’ capacity to bind Calcium.

**Fig 1.**
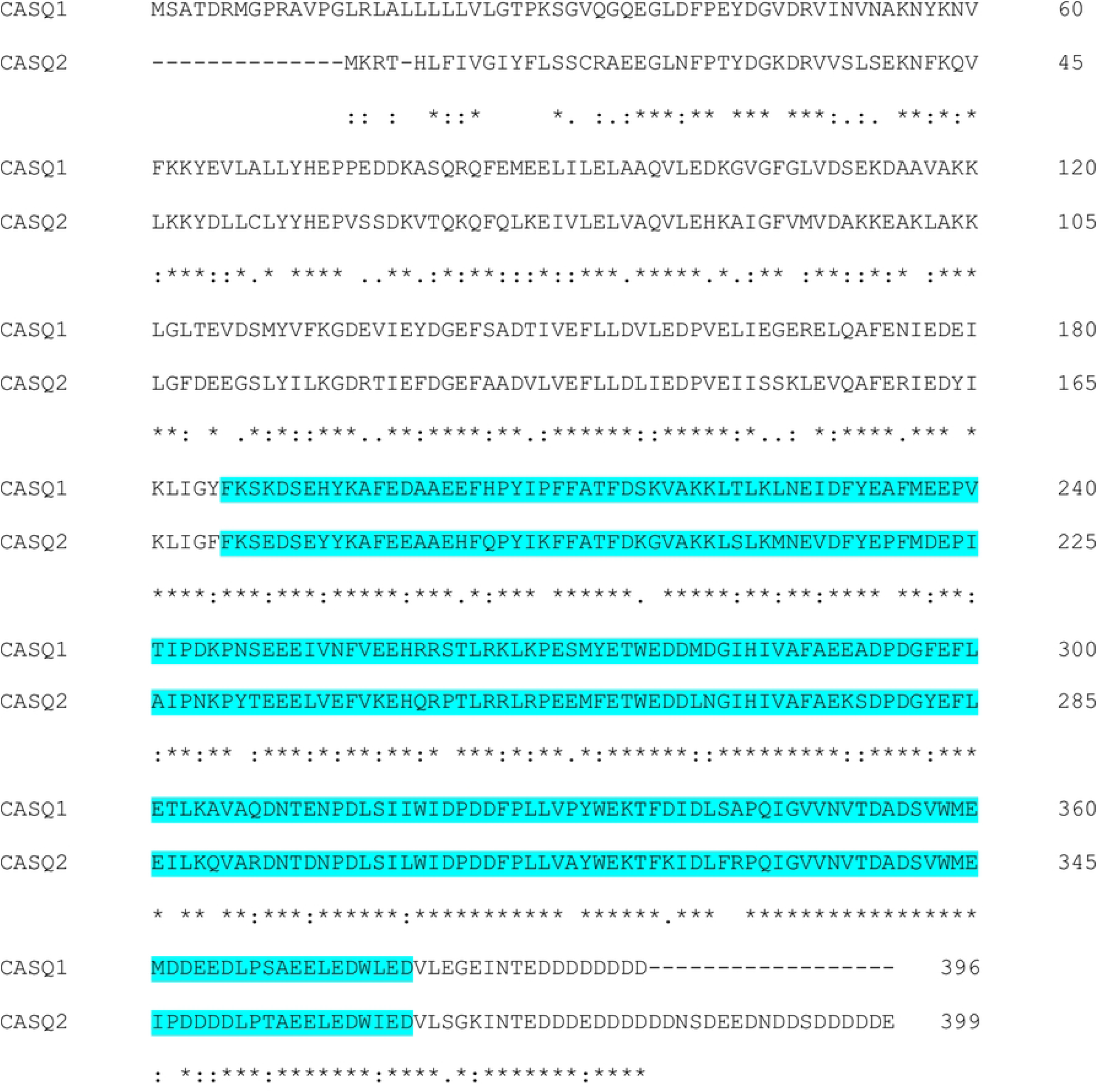
Alignment of CASQ1 and CASQ2 protein sequences. CASQ1 and CASQ2 protein sequences alignment shows 66.4% of identical residues (*). Thioredoxin6 predicted PFAM domain is highlighted in light blue for both proteins.

**Fig 2.**
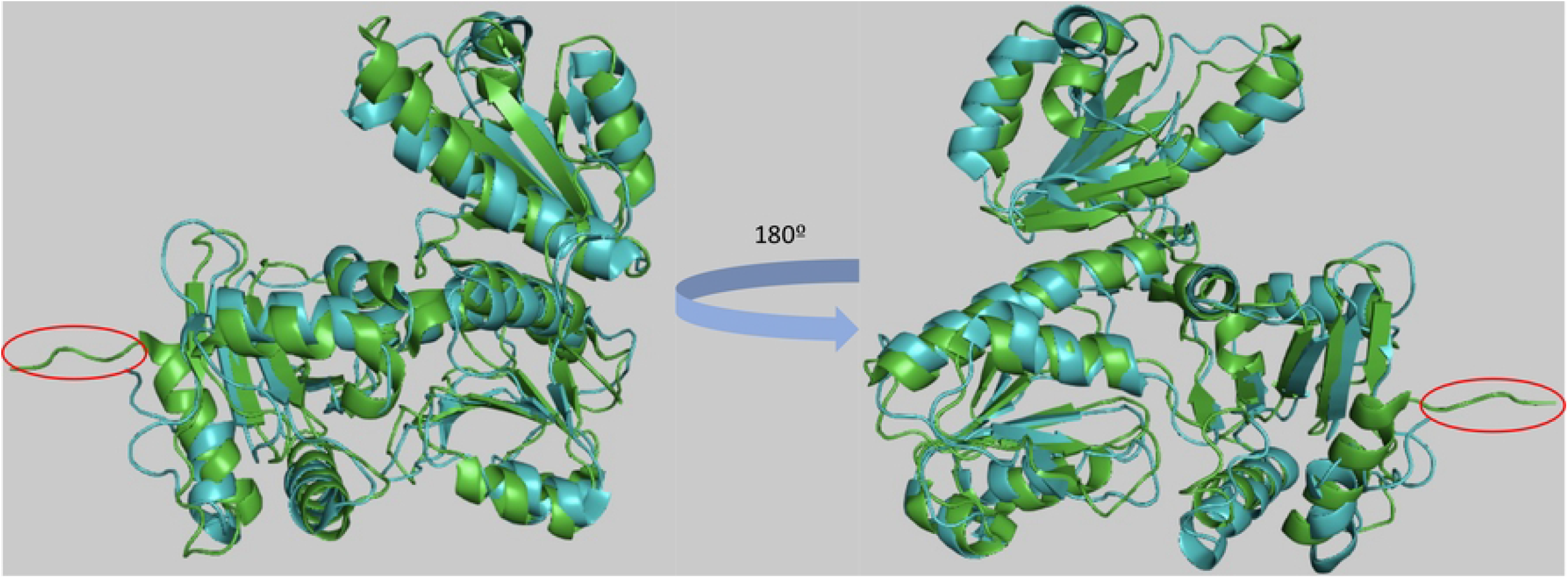
Superimposition of CASQ1 (PDB id:3UOM) and CASQ2 (PDB id:2VAF) protein structures. Solved 3D structure of CASQ1 and CASQ2 proteins are shown in superimposed cartoon representation. Conserved structure folding is shown between both proteins CASQ1 (Blue) and CASQ2 (Green), except for the extended C-terminal region of CASQ2, highlighted by a red circle.

### Gene expression analysis

Quantitative real-time PCR was performed to monitor the gene expression of *CASQ1, CASQ2, ICAM1, VCAM1*, and *SELE* in rat vessels at the site of injury. *GAPDH* was used as a reference gene. Gene expression was monitored at different timing post-injury (0, 30, 90 and 120 min). Untreated rat cDNA was used as a negative control for the relative expression of target genes. Results are expressed in fold change compared to the control group. Selected endothelial cells markers for inflammation, *VCAM1, ICAM1* and *SELE* were used to monitor the inflammatory process [32]. Figure 3 shows that *VCAM1* and *ICAM1* expression is upregulated starting from 30 min until 90 min post-injury as expected and reaching up to an 8-fold increase compared to the control group. The *SELE* expression profile shows a small delay in differential expression compared to CAMs; upregulation of expression was observed starting from 90 min (4-fold increase) and reaching a 7-fold increase at 120 min (Figure 3). *CASQ1* displayed no change in expression profile, however, interestingly for *CASQ2*, we observed a significant upregulated expression (*p*< 0.01) by more than 60-fold and 20-fold increase at 30 min and 90 min post-injury respectively (Figure 3), compared to the control group. The overall profile of the considered inflammatory markers is in line with our previous findings using the same animal model which delineates the importance of the *CASQ2* gene upregulation observation made.

**Fig 3.**
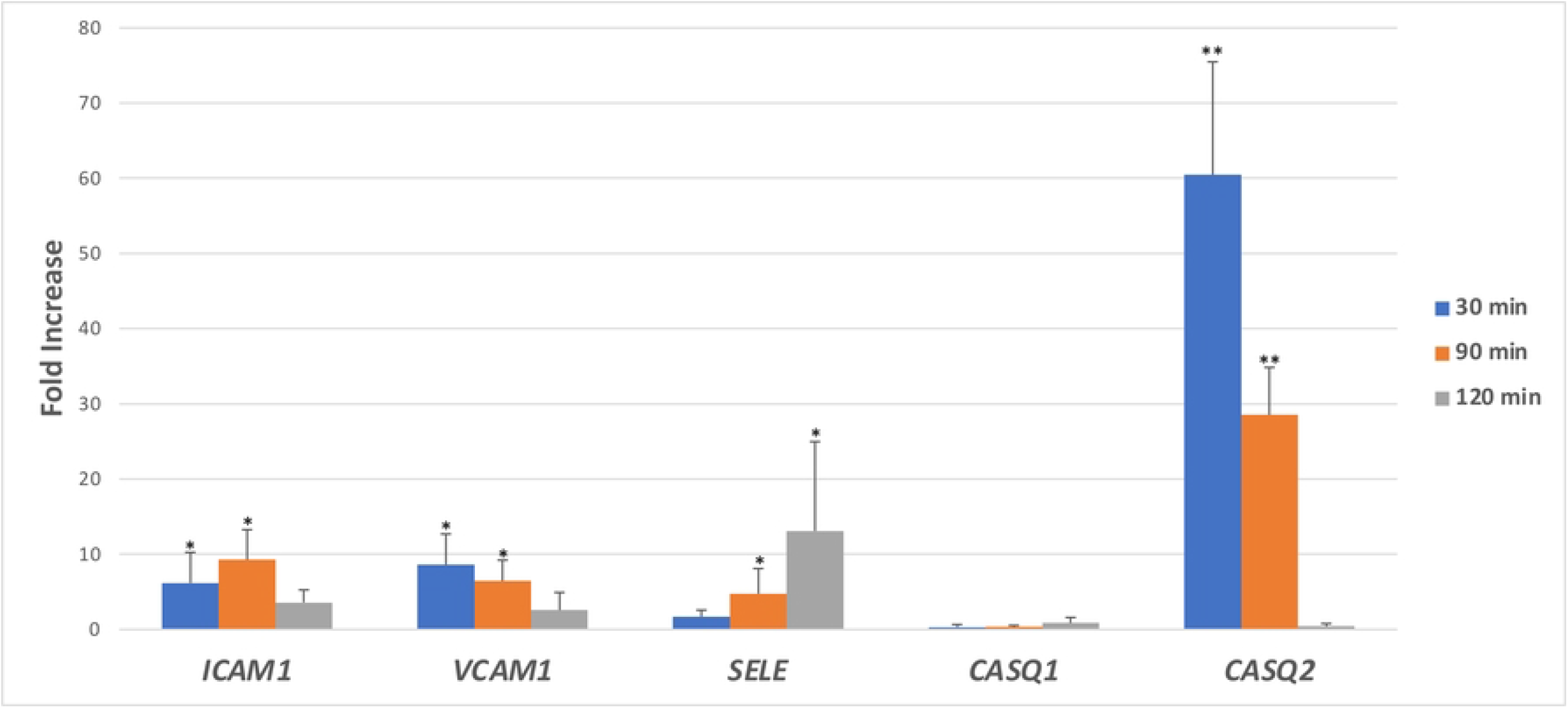
Gene expression profiling. *ICAM1, VCAM1, E-SELE, CASQ1* and *CASQ2* genes expression levels were monitored using real time PCR in vessels at the site of inflamed rat skeletal muscle at 30-, 90- and 120-min post-injury using the untreated control group as a reference (non represented). Results are expressed in fold change compared to the untreated control group (expression unit =1 for each time-point) and using GAPDH as a reference gene. (* *p*< 0.05 and ** *p*< 0.01).

### Multiplex Immunohistochemistry and Immunofluorescence

Histology of skeletal muscle injury in rat model by Hematoxylin and Eosin staining, showed that, as expected, an infiltration of neutrophils in the site of injury was observed 4 hours post-injury (Figure 4A and 4B). For *in situ* investigation, we performed a dual immunohistochemistry labeling of CASQ1 and CASQ2 that showed detectable expression of both proteins in healthy tissues, with a presence of CASQ2 mainly at the blood vessels in control sample tissues and CASQ1 in muscle tissues (Figure 5A). In the group of injured animals (Figure 5B), we observed a congestion in blood vessels at the site of injury together with infiltration of inflammatory cells in 30 minutes post-injury as reported previously [29]. This was concomitant with an upregulation of CASQ2 expression most likely in the smooth muscle wrapping the vessels present at the site of injury as shown by co-staining with CD34 (a specific marker of microvascular endothelial cells) in Figure 6 (multiplex Immunofluorescence) in comparison with the control group.

**Fig 4.**
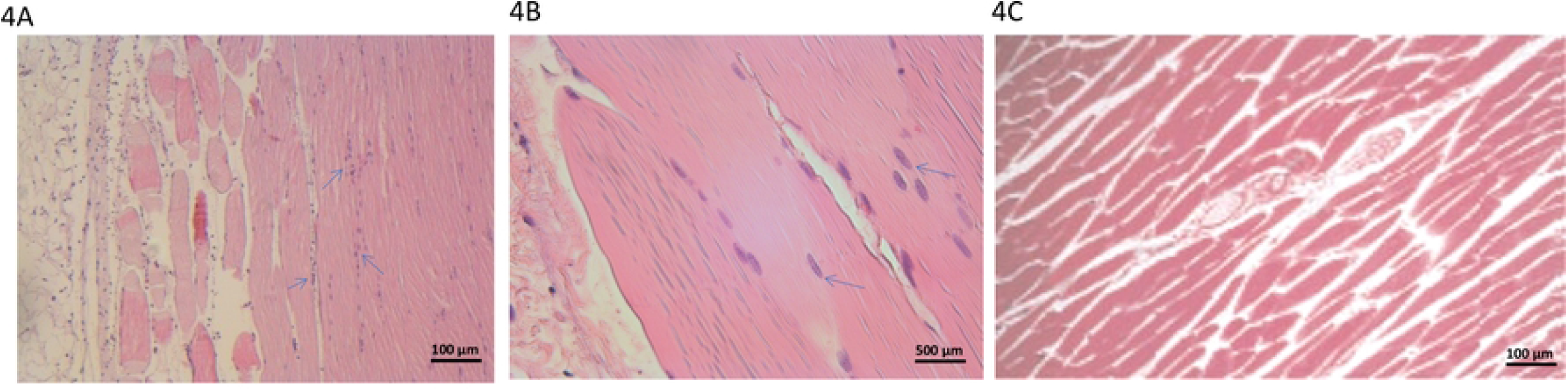
Histology of skeletal muscle injury in the rat model by Hematoxylin and Eosin staining. Images were acquired at 100X (A) and 400X (B) magnification showing neutrophils infiltration (blue arrows) in the site of injury at 4 hours post-injury in comparison to the non-injured control tissue (4C) acquired at 100X magnification.

**Fig 5.**
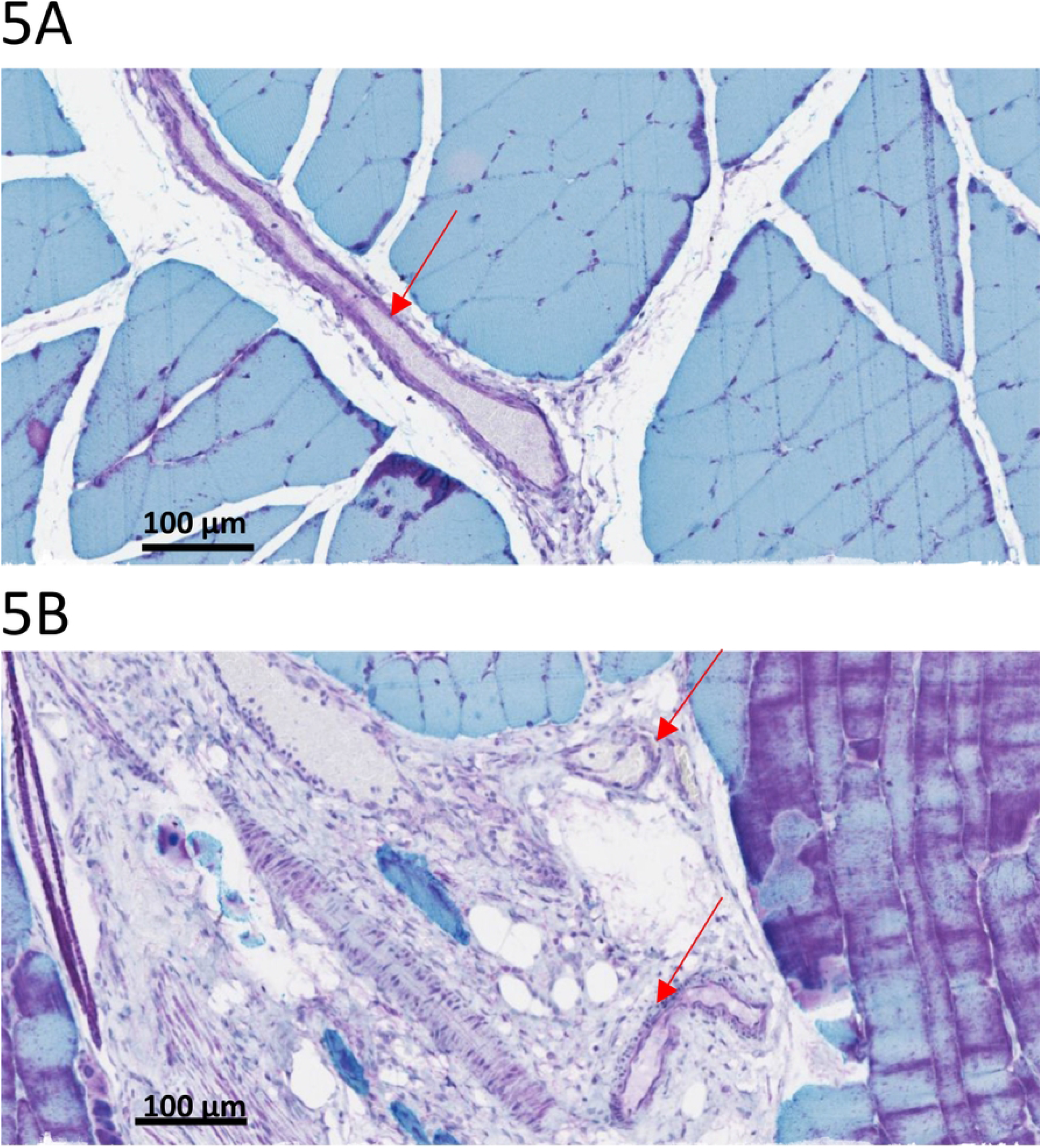
Multiplex Immunofluorescence labeling of CASQ1 (Teal) and CASQ2 (Purple) in healthy tissues (A) and 4-hours post-injury (B). Staining shows CASQ1 expressed mainly in muscular tissue and CASQ2 expressed mainly in vessels (indicated by red arrows).

**Fig 6.**
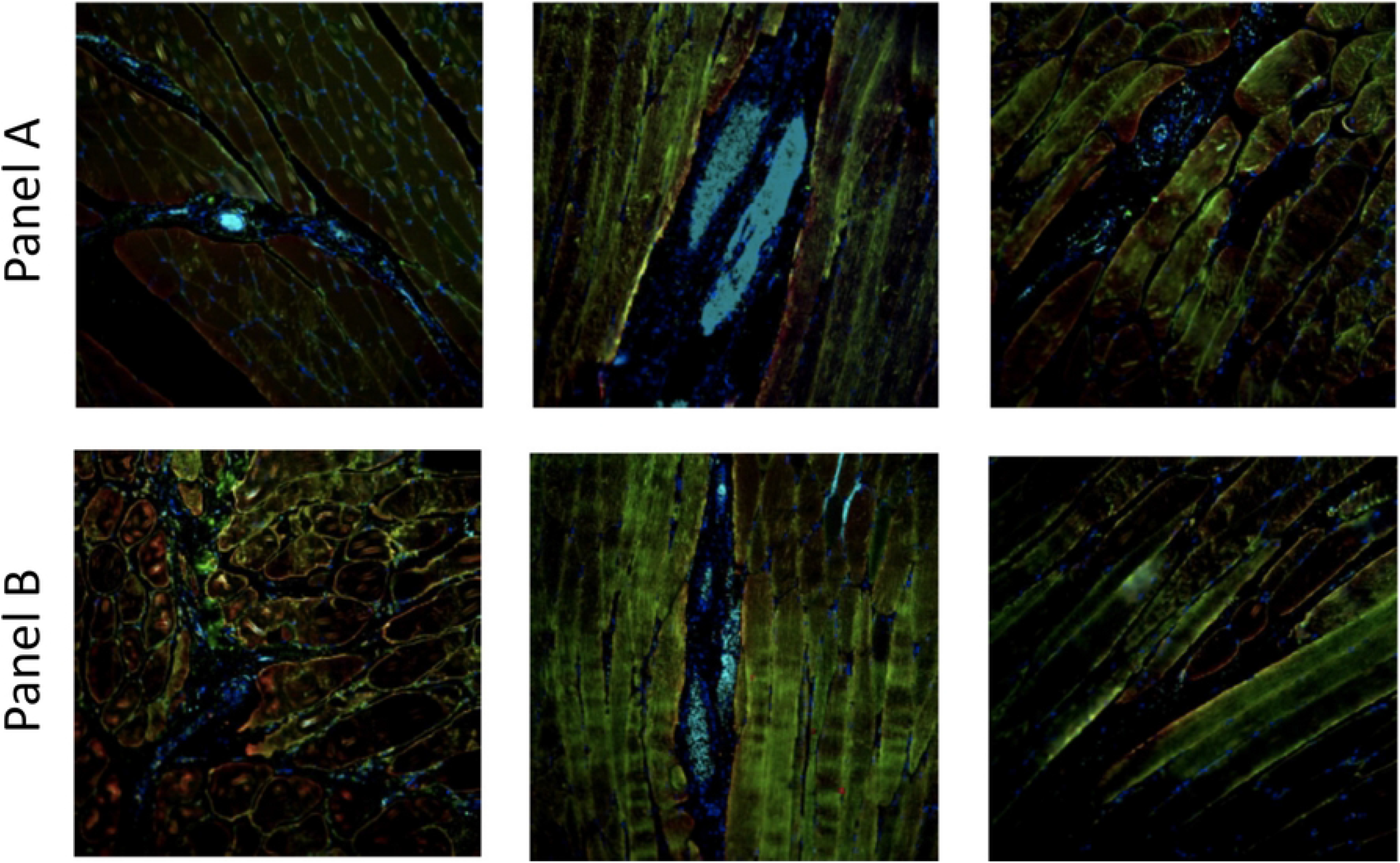
Multiplex Immunofluorescence staining of CASQ1(Red), CASQ2 (Green) and CD34 (Cyan) in healthy tissues (A) and 4-hours post-injury (B). DAPI (Blue), is used to label cell nucleus. Colocalization of CASQ2 and CD34 was detected in both panels A and B. Samples were collected from different animals of the study groups for each panel.

## Discussion

The function of the major mediators of inflammation such as the inflammasome and the leukocytes adhesions molecules is calcium dependent [33]. Therefore, availability of the necessary amount of calcium and its efficient mobilization are important for the inflammatory process to develop, in infections, sepsis, autoinflammatory diseases, and metabolic diseases and trauma (Torre-Minguela et al. 2017). In acute inflammatory diseases, the mediators of inflammation are timely overexpressed in the inflamed tissues and in the vascular compartment while in chronic inflammatory states the expression of these mediators is steady. Consequently, in both situation the amount of readily available calcium should determine the right extent of the inflammatory process. In this study we show that CASQ2 or Calsequestrin isomer 2, a member of the thioredoxin protein family is significantly overexpressed in inflamed mechanically injured rat skeletal muscle at the level of congested vessels. This overexpression occurs between 30 and 90 minutes post injury which is concomitant with leukocytes infiltration in the injured skeletal muscle (first wave of neutrophils transmigration one hour post-injury) as previously reported in the rat model of skeletal muscle injury [29] and observed in this study. This observation is consistent with the fact that the adhesion molecules that mediate leukocytes transmigration into inflamed tissues depends heavily on the mobilization of Ca^2+^ [34]. Interestingly, Calsequestrin is the major Ca^2+^ binding protein and the main Ca^2+^ storage and buffering protein [18]. Indeed, CASQ is present in the sarcoplasmic reticulum, and is an important regulator of Ca^2+^ release channels in both skeletal and cardiac muscle [18]. In addition, this study suggests that CASQ2 overexpression occurs in the smooth muscle that wraps the vessels present at the site of injury as shown by immunohistochemistry and immunofluorescence. Indeed, while the expression of CASQ1 is well-established in skeletal muscle, CASQ2 has been shown to be expressed in smooth muscle [18]. Volpe et al. have demonstrated the existence in smooth-muscle cells of discrete endoplasmic-reticulum areas specialized in the rapidly exchanging Ca2+ storage and release [35], and thus in the control of a variety of functions, including smooth-muscle contraction. Our observation of a marked overexpression of the CASQ2 isomer in the vessels of smooth muscle at the site of an inflammatory injury suggests that in addition to being an essential component of smooth muscle excitation-contraction, we suggest that CASQ2 provides high local concentration and release of the Ca^2+^ necessary to activate the overexpressed leukocytes and endothelial adhesion molecules (ICAM1, VCAM1 and Selectin) that promote the adhesion to vascular endothelium and leukocytes over flux into the inflammation site as early as 30 min post-injury. In addition, smooth muscle cells contribute or regulate the inflammatory reaction in the progression of atherosclerosis, especially in the context of the activation of various membrane receptors, and how they may regulate vascular inflammation [36, 37].

Hübner and collaborators [13] demonstrated that for neutrophils, the calcium dynamics of an individual cell correlates with its respective functional state and that neutrophils specific calcium dynamics carries crucial information on their functions and phenotype [13]. Hence our finding is limited to the concomitant overexpression of CASQ2 with the over flux of neutrophils at the inflammatory injury site suggests CASQ2 as a potential mediator of this calcium dynamics, and further experiments involving CASQ2 knock-out animal model together with calcium dynamics monitoring wuld help confirm our observations.. Although the thioredoxin family is a multi-functions protein family [38] endowed mainly with multiple protein chaperone functions, to our best knowledge CASQ proteins are not involved in such functions. The main role of these proteins is the storage and release of calcium ions. We speculate that CASQ2 overexpression in the smooth muscle of inflamed vessels but not CASQ1 could be linked to the difference in calcium binding affinity of the two proteins and the tissue specific expression [39], allowing for modulation of calcium influx.

To our best knowledge, This is the first observation of CASQ2 overexpression in congested vessels within a mechanically injured skeletal muscle. Therefore, we e propose CASQ2 as a calcium source for the inflammasome and hence a putative biomarker for muscle inflammation. Further analysis of CASQ2 knock-out mice model will help understanding the exact role-played by this protein in inflammation.

## Acknowledgements

We thank Mr. Ammar Marweni from the Arabian Gulf University for his support in the work involving animals.

## Ethical approval

Experimental protocols were approved by the Arabian Gulf University Research and Ethics committee. All methods were carried out in accordance with the committee’s relevant guidelines and regulation.

